# Prediction errors indexed by the P3 track the updating of complex long-term memory schemas

**DOI:** 10.1101/805887

**Authors:** Franziska R. Richter

**Affiliations:** Cognitive Psychology Unit, Institute of Psychology, Leiden University, Leiden, The Netherlands

**Keywords:** schema, long-term memory, ERP, P3, P300, prediction error, updating, knowledge

## Abstract

Memory schemas are higher-level knowledge structures that store an abstraction of multiple previous experiences. They allow us to retain a multitude of information without the cost of storing every detail. Schemas are believed to be relatively stable, but occasionally have to be updated to remain useful in the face of changing environmental conditions. Once a schema is consolidated, schema updating has been proposed to be the result of a prediction-error (PE) based learning mechanism, similar to the updating of less complex knowledge. However, for schema memory this hypothesis has been difficult to test because no sufficiently sensitive tools to track modifications to complex memory schemas existed so far. Current research on the updating of less complex beliefs and at much shorter time scales has identified the P3 as an electrophysiological correlate of PE-induced updating of beliefs. In this study, I recorded electroencephalography and continuous memory measures during the encoding of schema consistent vs. inconsistent material to test the behavioural and neural correlates of schema updating. I observed that PEs predicted the updating of a schema after a 24-hour delay, especially when participants were faced with inconsistent compared to consistent material. Moreover, the P3 amplitude tracked both the PE at the time of learning as well as the updating of the memory schema in the inconsistent condition. These results demonstrate that schema updating in the face of inconsistent information is driven by PE-based learning, and that similar neural mechanisms underlie the updating of consolidated long-term memory schemas and short-term belief structures.

## Introduction

Memory schemas are complex dynamic knowledge structures that allow us to store vast amounts of information in a concise manner. Schemas evolve slowly over time, and are generally believed to be stable once consolidated (Ghosh & Gilboa, 2014). However, in a dynamically changing world, schemas need to be adjusted or updated from time to time (Piaget, 1952). *Accommodation* is a hypothesized updating mechanism that has been suggested to lead to small changes to the schema’s structure itself, in response to changing environmental conditions (Ghosh & Gilboa, 2014).

One popular idea of how memory schemas can be updated or made to *accommodate* new information is by means of prediction-error (PE) based learning (cf. Henson & Gagnepain, 2010; van Kesteren, Ruiter, Fernández, & Henson, 2012): When a predicted outcome does not match our experience, the resulting PE provides us with a surprise signal that is believed to guide learning. Recent findings suggest that PE-based learning underlies memory updating in many contexts: the updating of rules (Greve, Cooper, Tibon, & Henson, 2019), of episodic memories (Greve, Cooper, Kaula, Anderson, & Henson, 2017; Sinclair & Barense, 2018, 2019), and of single semantic facts (Pine, Sadeh, Ben-Yakov, Dudai, & Mendelsohn, 2018). However, research on the accommodation of new information into complex abstracted knowledge structures such as memory schemas is scarce.

In the context of *short*-*term* decision-making tasks the neural mechanisms of PE-based updating have been studied with functional magnetic resonance imaging (e.g., Behrens, Woolrich, Walton, & Rushworth, 2007) and electroencephalogram (EEG) (Bennett, Murawski, & Bode, 2015; Jepma et al., 2016; Kolossa, Kopp, & Fingscheidt, 2015). In these tasks participants adjust their predictions multiple times within an experimental session. Such PE based-updating of short-lived beliefs has been shown to be indexed by the P3 (or P300) signal (Bennett et al., 2015; Jepma et al., 2018, 2016; Kolossa et al., 2015). The P3 is an event-related potential (ERP) that has been linked to updating of the (immediate) context in a number of tasks that require the comparison of a current stimulus with a recently preceding stimulus (Polich, 2012), for example in oddball and working memory tasks (Donchin, 1981; Sutton, Braren, Zubin, & John, 1965). This ERP has been suggested to be divisible into two subcomponents, the fronto-centrally distributed P3a and the P3b, which has a centro-parietal topography (see Fonken, Kam, & Knight, 2019, for a review). The more frontally distributed P3a has been linked to mismatch and novelty processing, and shown to be dependent on the hippocampus, while the P3b, a parietally centred component, is more strongly linked to target detection, and possibly not as strongly linked to hippocampal processing (Fonken et al., 2019). In the current study, I investigated both P3a and P3b to assess which of the two ERPs, or both, or any, correlated with PE size and schema updating, based on prior findings of both P3a (Bennett et al., 2015; Kolossa et al., 2015) and P3b (Jepma et al., 2018, 2016) involvement in short-term belief updating tasks.

Of note, belief updating in the decision-making experiments mentioned above takes place on a much smaller time scale than what is thought to be necessary for the formation and updating of memory schemas. These much more complex knowledge structures are assumed to be extracted over hours, days, or even months (Ghosh & Gilboa, 2014; O’Reilly, Bhattacharyya, Howard, & Ketz, 2014). While short-lived believes about the current situation and schemas that have been built up over prolonged periods of time share many conceptual similarities (e.g., extraction of generalities, acceptance of variability in the data) it is currently unknown whether the same neural mechanisms underlie the updating of short-lived beliefs and memory schemas. Furthermore, how long-lasting the effects induced by a putative updating of a schema are, is also unclear.

An argument for similar neural mechanisms in the updating of short-lived beliefs and complex memory schemas is in line with behavioural evidence which suggests that PE-based learning also works on longer time-scales (e.g., Greve et al., 2019; Pine et al., 2018). Furthermore, initial evidence suggest that for some types of memory, such as semantic knowledge, updating relies on similar regions of the mid-brain that are also thought to underlie the updating of more short-lived beliefs (Garrison, Erdeniz, & Done, 2013; Pine et al., 2018). In addition to updating, the P3a and the P3b have been linked to better encoding in general, such that larger amplitudes of either potential are associated with better memory in a subsequent test (Azizian & Polich, 2007; Knight & Scabini, 1998; Richter & Yeung, 2016). The link between P3 and subsequent memory could indicate that PE-induced updating effects will have prolonged consequences rather than only affecting behaviour in the short term.

It is therefore conceivable that the P3 might also index updating of long-term memory schemas and that these effects persist over long periods of time. A consolidated schema should allow for mismatches (PEs) between schema-based predictions and new inconsistent information. These PEs should consequently induce small modifications to a schema (Ghosh & Gilboa, 2014; cf. Piaget, 1952). With regards to schema memory, however, a major difficulty lies in the tracking of such subtle changes to these complex knowledge structures. Small changes in memory schemas have only recently been made visible in humans using sensitive continuous memory tools (Richter, Bays, Jeyarathnarajah, & Simons, 2019) (see Richards et al., 2014 for an example in the animal literature). This research has made updating processes visible and now allows studying the neural correlates of schema updating. It also suggests that the original schema can have long-lasting effects on later behaviour.

In this study, I set out to test the putative (neural) mechanisms underlying schema updating: PE-based learning processes indexed by the P3. To do so, I am employing a continuous memory task to study the neural mechanisms of schema updating using EEG. By using a prediction-based learning paradigm, I am able to track the neural correlates of trial-by-trial prediction violations and relate them to schema updating one day later. Based on the findings reviewed above, I predict that the P3 signal should track prediction violations and updating mechanisms in the context of memory schemas.

## Materials and Methods

### Materials

The stimulus set used in the current study consisted of 400 images of 4 different categories: animals, clothes, furniture, and food (fruits and vegetables), and largely overlapped with those used in a recent study on schema memory (Richter et al., 2019). Backgrounds were removed for each object picture and stimuli were presented in a size of ∼150 × 150 pixels on top of a 400 × 400 pixel background image. The background image was used on every trial throughout the experiment to facilitate encoding of the locations. On this background image the location of the pictures was restricted to locations on an invisible circle, consistent with the approach used in other paradigms studying memory precision for location information (Cooper et al., 2017; Harlow & Donaldson, 2013; Harlow & Yonelinas, 2016; Murray, Howie, & Donaldson, 2015; Richter, 2020; Richter et al., 2019; Richter, Cooper, Bays, & Simons, 2016). As outlined in more detail below, each category was predominantly associated with specific circle locations and participants had to learn these ‘schemas’.

### Procedural overview

Twenty-five participants took part in the current study. Participants were reimbursed for their participation with payment or course credits. After reading an information sheet on the experiment and giving informed consent according to the declaration of Helsinki, they were presented with task instructions.

A firmly established schema is the foundation for schema-based prediction violations to occur, which is to say that schema updating requires a schema to be at least partially consolidated. To achieve this participants came to the lab on three consecutive days, with sessions scheduled 24 hours apart (see Figure 1 for an overview). Participants in the current study first acquired location schemas for the above mentioned stimulus categories in a prediction task on day 1. On day 2 they experienced inconsistencies in a subset of the categories while their EEG was recorded. Later, on day 3 they were tested on their memory for all stimuli. Continuous memory precision measures were used to track the formation and updating of memory schemas, similar to an approach used recently in a behavioural paradigm (Richter et al., 2019).

**Figure 1.**
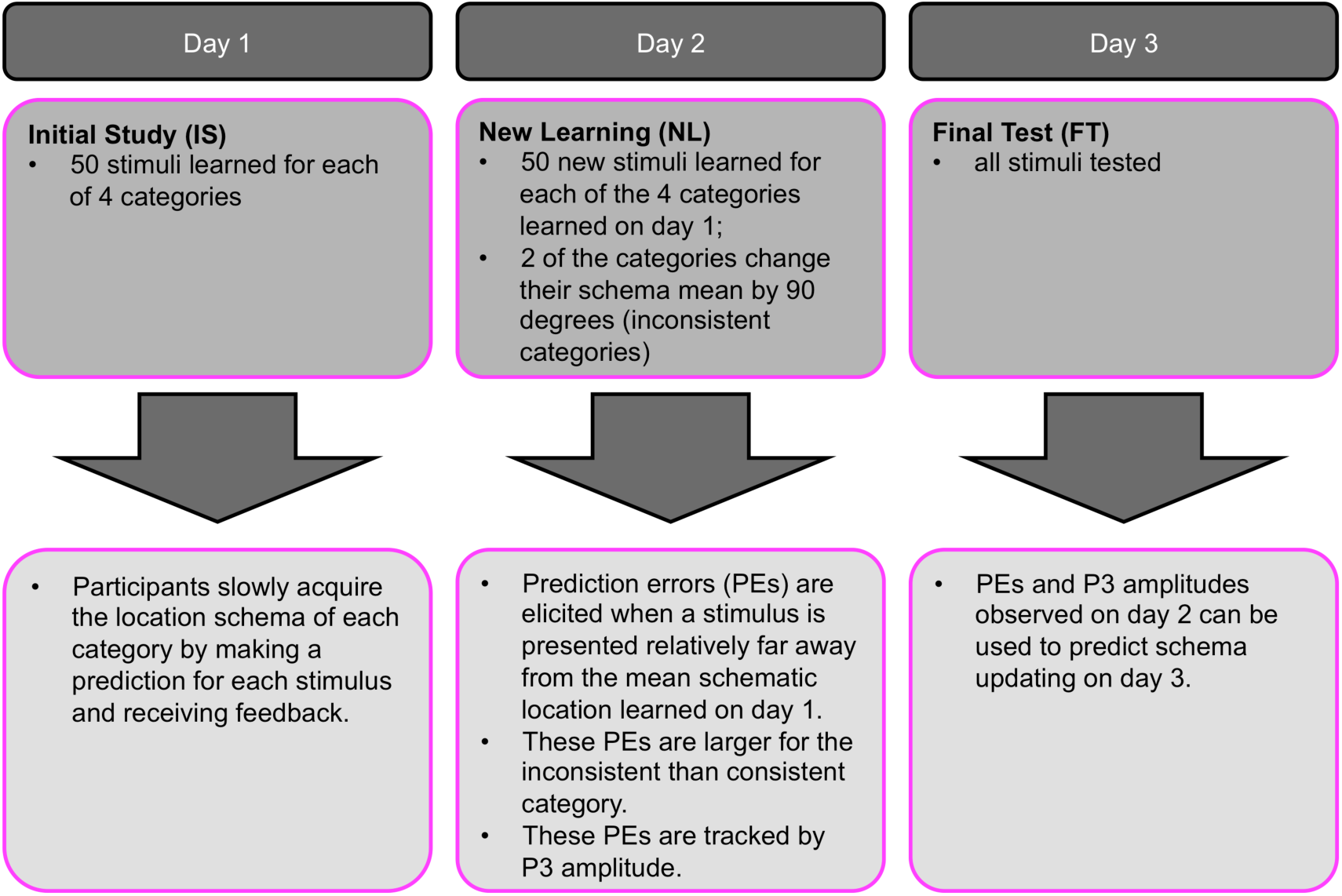
Sessions of the main experiment and expected behavioural effects. On day one participants learn stimuli in each category by performing a prediction task, and acquire location schemas. On day 2 new stimuli of each of the categories are learned, but two of the categories become inconsistent, inducing PEs evident in behaviour as well as increased P3 amplitudes. On the final day, the participants complete a precision memory test for all items. It is hypothesized that behaviour in this day 3 test can be predicted from day 2 behavioural and EEG measures.

I assessed the hypotheses that introducing information that is inconsistent with a previously learned memory schema 24 h after the initial schema had been acquired, would lead to an increase in PEs, relative to a consistent condition, in which newly learned information was congruent with a previously learned schema. Furthermore, I predicted that these PEs would be accompanied by increased P3 amplitudes and that the size of these two measures, PEs and P3 amplitudes, would predict the degree to which a schema is updated on day 3.

On day 1, participants first completed a practice task (34 trials) using pictures of musical instruments that did not overlap with the image categories used in the main experiment. The 34 practice trials were spread over 4 blocks with a length of 2, 4, 10, and 18 trials, to give participants a chance to get used to the prediction task and ask the experimenter additional questions after the blocks. The participants’ task, which is described in more detail below, was to predict where on an invisible circle a picture should be placed. They subsequently had to encode the correct location that was provided as feedback after their prediction. In the practice phase, most (76.47%) of these correct locations, that is, the items’ feedback locations presented to participants, occurred in a 90-degree segment centred on the 65 degree point of the circle. Because participants did not know about this pattern, they initially guessed where a certain musical instrument picture would be placed until they learned that stimuli followed a location schema, and that instruments would ‘cluster’ around this specific 65 degree point on the invisible circle.

In the main task participants learned stimuli from all 4 categories on day 1 and 2. The goal of day 1 was for participants to learn that most of the stimuli in each of the 4 categories were presented in category-specific areas of the screen. For example, animals would be clustered in a 90-degree window centred at 110 degrees of the circle (see Figure 2A). The mean locations, in degrees, for the four schemas were 20, 110, 200, and 290 degrees. The assignment of categories to these mean locations – or *schema means* – was counterbalanced across participants. Within each category most trials (40 out of 50, *inside quadrant trials*) fell within a category specific 90-degree window. To make the location schemas less obvious and ensure gradual learning 10 out of 50 trials fell outside of this 90-degree window (*outside quadrant trials*). In an initial study phase on day 1 and during a new learning phase on day 2, participants completed 10 blocks of 20 trials of schema learning with 50 trials each for each of the four categories. Within the 10 blocks, stimuli of all 4 categories (animals, clothes, food, and furniture) would be presented in equal proportions.

**Figure 2.**
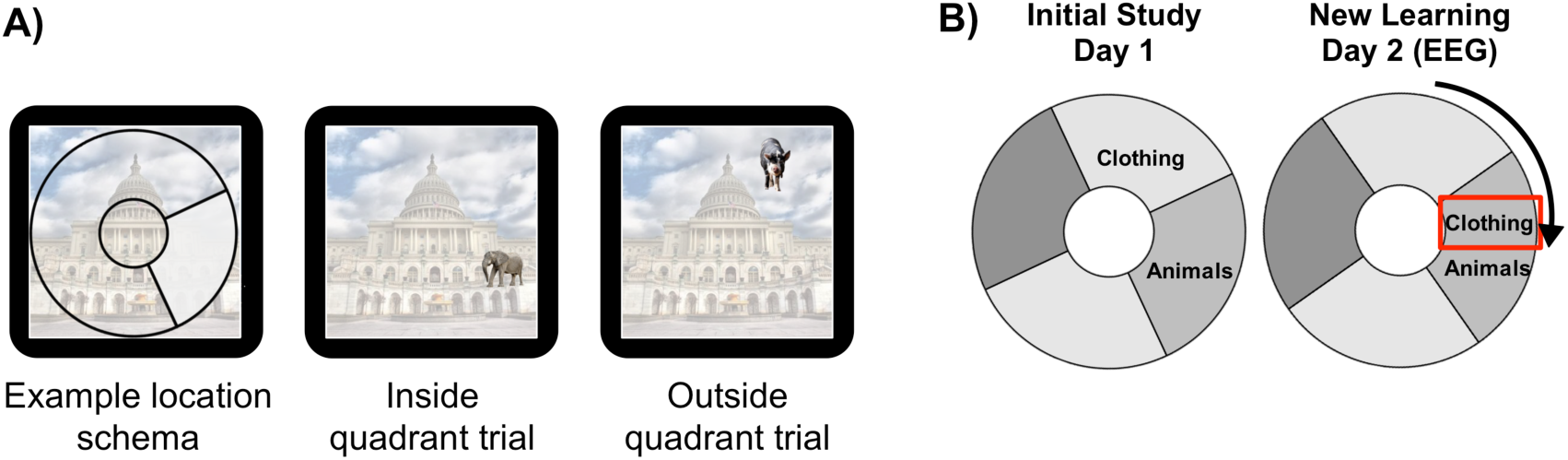
Location schema for the animal category. **(A)** Illustration of the size of the circle segment occupied by a category, in this case animals. In *inside quadrant trials* the images of the category are shown within the 90-degree circle segment, in *outside quadrant trials* they are presented outside of it. **(B)** On day 2 the category-specific circle segment changes for two of the four categories. In this example the clothing category is moved by 90 degrees clockwise.

On day 2 the assignment of categories to circle locations was retained for 2 of the 4 categories, but was changed for the other 2. For these ‘inconsistent’ categories the schema mean would shift by 90 degrees clockwise (see Figure 2B). For example, if the schema mean for the animal category was 110 degrees on day 1, it would be 200 degrees on day 2. The 200 stimuli (50 per category) learned on day 2 did not overlap with those learned on day 1 as they were new exemplars of the same categories studied on day 1. Which category was presented at which location was counterbalanced across participants, and the category shift always occurred in such a manner that all categories would be centred either around 20 or around 200 degrees. As a result, the consistent and inconsistent categories were matched for their visual location.

On day 3 participants completed a final precision test on all stimuli learned on the previous 2 days. They were presented with a previously studied stimulus at a random location and had to recreate the ‘correct’ location from memory. That is, they had to recreate the location that was provided to them as feedback for this stimulus in each trial on day 1 or day 2.

### Task

Trials on day 1 and 2 consisted of the following steps: First an objet was presented to participants at a random location on an invisible circle centred on a background image (see Figure 3A) with a white fixation cross presented in the centre of the screen. Participants had 5 seconds to move the picture to a position of their choice using the “g” (counter-clockwise) and “h” (clockwise) keys on a standard computer keyboard. They were instructed to use the space bar to lock their response. At first, participants would guess the location, as no prior schema was established yet. They subsequently had 3 seconds to rate the confidence in their choice on a 100 point scale ranging from certainly incorrect to certainly correct. They again used the “g” (left) and “h” (right) keys to move the slider and locked the response using the space bar. When 1 second was left for participants to make their decision, the fixation cross in the centre of the screen or the tick mark on the confidence scale would turn red to remind participants to lock their answer. If participants did not manage to lock their answers in time to confirm their response, the last location the participant moved the cursor to in the confidence rating or the location of the picture was recorded as the response. After this step, the picture was presented in the ‘correct’ location for 1 second (feedback). The picture then disappeared from the background leaving only a fixation cross and participants had another 4 seconds to memorize the seen location before the next trial started.

**Figure 3.**
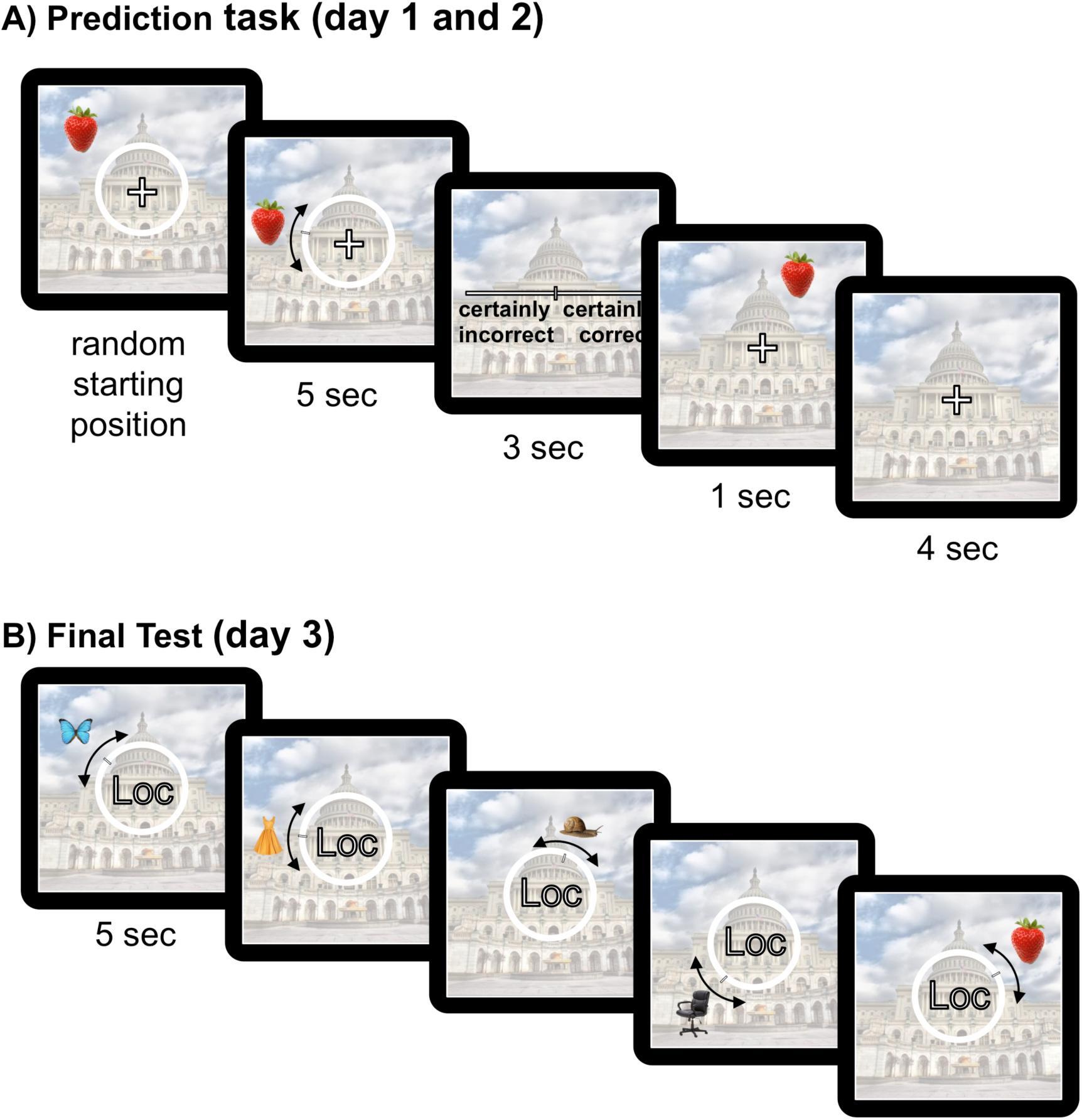
Task structure. (**A**) Prediction task on day 1 and 2. Participants are presented with an object at a random location and then have 5 seconds to move this object to a location of their choice. They subsequently have 3 seconds to indicate their confidence in their choice. The correct location (feedback) is presented to them for 1 seconds and participants then have another 4 seconds to memorize this location before the next trial starts. (**B**) In the memory test on day 3 participants are presented with the previously studied objects of all categories in a random order. Pictures are presented at a random location initially, and participants have to recreate the correct location that they have learned on day 1 or 2. Images can be moved around the entire circle in continuous steps. Loc = In the experiment the word ‘Location’ was presented instead of the abbreviation ‘Loc’.

The task on day 3 was slightly different from that on day 1 and 2. Participants were presented with all of the stimuli learned on days 1 and 2 in a random order. A stimulus was presented at a random location, with the word ‘Location’ presented in the center of the screen in white font. Participants were instructed to move the stimulus to the ‘correct’ location, that is the location that they were presented with in the feedback period at the end of each trial on day 1 and day 2. When 1 second was left for participants to make their decision, the word location would turn red to remind participants to lock their answer. If participants did not lock in their answers in time to confirm their response, the last location of the picture was recorded. Stimulus presentation was implemented using Matlab (Mathworks, Natick, MA) using Psychtoolbox (www.psychtoolbox.org) software.

### EEG data recording and analysis

The EEG was recorded from 32 scalp electrodes on a Biosemi system using AgCl electrodes. Electrode location corresponded to FP1, FPZ, FP2, F7, F3, FZ, F4, F8, FT7, FC3, FCZ, FC4, FT8, T7, C3, CZ, C4, T8, TP7, CP3, CPZ, CP4, TP8, P7, P3, PZ, P4, P8, POZ, O1, OZ, and O2. Two electrodes were attached on the outer canthi of both eyes, and two further electrodes were attached above and below the left eye to record blinks and eye movements. Left and right mastoids were also recorded. A sampling rate of 512 Hz was used for recording. Before analysis, the EEG data were downsampled to 100 Hz. A common mode sense electrode was used for online referencing and electrodes were re-referenced offline to the average of the two mastoids to remove any lateral biases.

Data processing was done separately for each participant. Raw EEG data were high-pass filtered at 0.5 Hz to remove drift. A low-pass filter of 40 Hz was used for analysis using the “eegfiltnew” method implemented in the EEGLab toolbox. Data were epoched into trials of 4500-ms duration to avoid edge artefacts in further analyses. This time-window included a 1500-ms pre-event period. Preliminary artefact rejection was done prior to independent component analysis (ICA) by removing trials with extreme amplitudes (±2000 µV across the entire epoch, as well as ±100 µV in a 100 ms baseline period). Any trial beyond 5 standard deviations in a single channel or all channels was considered improbable data and was furthermore rejected using EEGLab’s algorithm.

Data were baseline corrected in the time period −100 ms to 0 ms. Ocular artefact reduction was performed using ICA in EEGLab to identify blink components, which were mathematically subtracted from the data. In a final artifact rejection step following the removal of the ICA components, trials were rejected if at least one electrode showed a difference of more than ±75 µV in a time window from the 100 ms baseline period to 900 ms after the event, and accordingly within a total period of 1 second.

All further data analyses were completed in MATLAB using custom-written routines. The lowest number of trials included for a participant after artifact correction was 111 out of 200 trials. All other participants had more than 122 trials. No participant was excluded. Further data analysis steps are described for the relevant analyses below. In the analyses below violations of Sphericity were assessed. If the Sphericity assumption was violated corrected *p*-values according to Greenhouse-Geisser will be reported alongside *original* degrees of freedom to maintain readability.

Due to a technical error, data from 1 block on day 1 was lost from one participant, so this participant’s data was not included in the day 1 analyses. When reporting results form day 2 and 3, I averaged across the two consistent categories (consistent condition) and the two inconsistent categories (inconsistent condition) in all analyses reported below.

## Results

### Behavioural Results

#### Schema Learning on Day 1 and Day 2

To investigate whether participants learned the location schemas, I tracked their absolute error in the prediction task on day 1. Of note, since the pictures were uniformly distributed within a category’s 90-degree segment, participants could not reduce their error beyond a certain point, even if they learned the schema perfectly. As evident from Figure 4, participants’ performance in the task improved over the course of the initial study phase, which occurred on day 1. Performance differed significantly across blocks, *F*(9, 207) = 9.726, *p* < .001, Sphericity not assumed. Follow-up polynomial contrasts indicated a significant linear trend, *F*(1, 23) = 35.664, *p* < .001, indicating a decrease in the absolute error across blocks on day 1. Hence, participants learned the schema over the course of day 1, evident in a decreasing PE.

**Figure 4.**
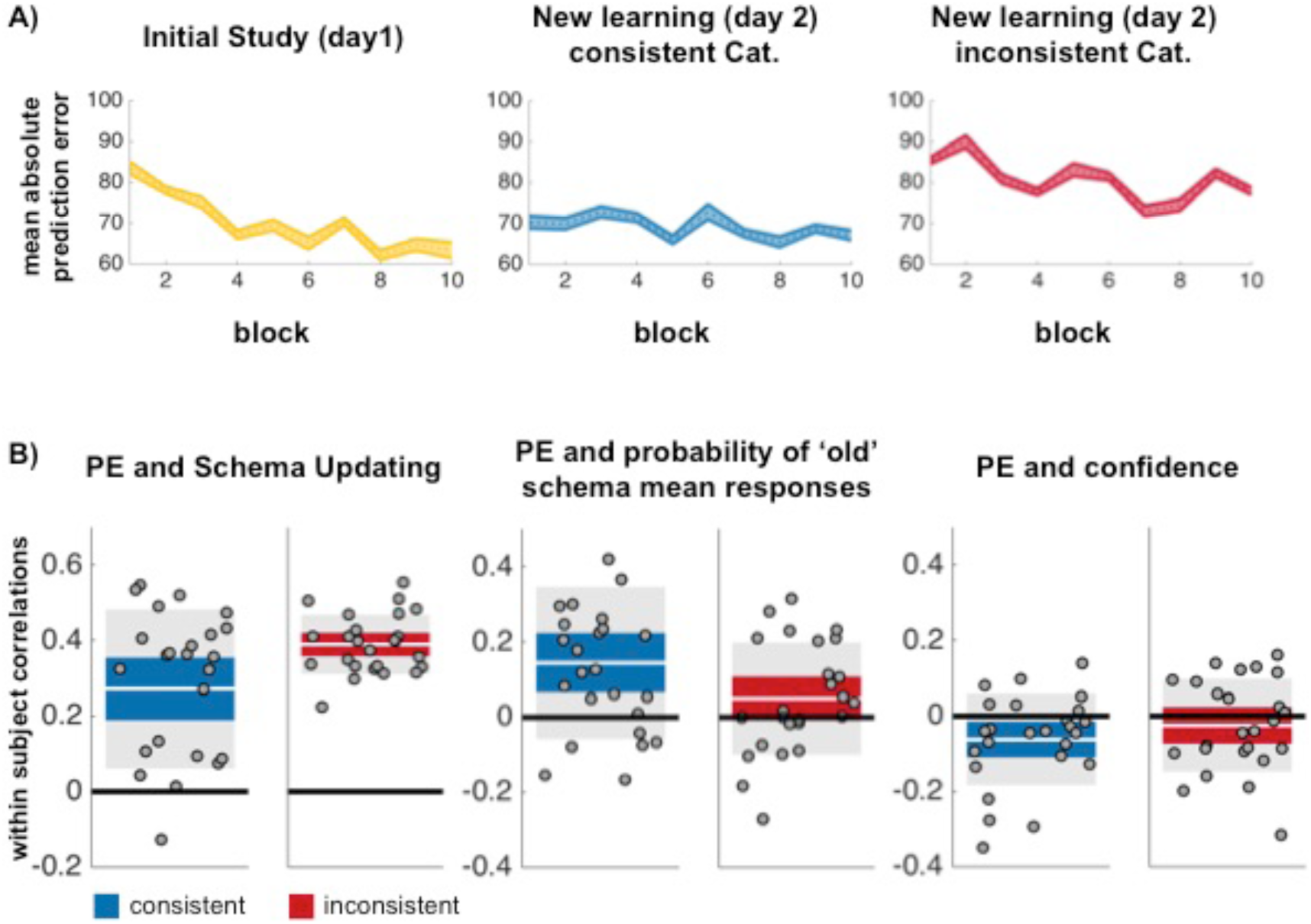
**(A)** Mean absolute PE (dotted line) across individual blocks of initial study (yellow) and new learning. For the new learning phase performance is plotted separately for the consistent categories in blue and the inconsistent categories in red. Solid lines reflect standard errors of the mean. **(B)** Within-subject correlations between PE and schema-updating score (left), between PE and the probability of responses consistent with the *old* schema mean according to the computational model (center), and between PE and confidence (right). In each plot the consistent condition is displayed in blue on the left and the inconsistent condition in red on the right. Each circle represents one subject. The white line represents the mean and the blue/red area the 95% confidence interval. The grey areas indicate *mean* ± 1 *SD.* Illustration of these correlations was created via the notBoxPlot function in Matlab https://nl.mathworks.com/matlabcentral/fileexchange/26508-notboxplot.

On day 2 participants’ responses should be guided by this schema in the consistent categories. In the inconsistent categories, in which the mean location for the category shifts by 90 degrees, participants should update their schema after an initial increase in PEs. To test this prediction, I first calculated an ANOVA with factors consistency (consistent and inconsistent) and block (1 to 10). The ANOVA revealed a significant main effect of consistency, with lower errors in the consistent conditions, *F*(1, 24) = 14.095, *p* = .001, Sphericity assumed. In addition, a significant main effect of block was observed, *F*(9, 216) = 2.589, *p* = .007, Sphericity assumed, as well as a marginal interaction between consistency and block, *F*(9, 216) = 1.737, *p* = .082, Sphericity assumed.

Due to the a priori hypothesis that PE on day 2 should decrease from block 1 to block 10 in the inconsistent but not, or not as strongly, in the consistent categories, I conducted several follow-up tests. Consistent with expectations, there was no significant main effect of block, *F*(9, 216) = 0.945, *p* = .487, Sphericity assumed, and no significant linear trend of block, *F*(1, 24) = 1.512, *p* = .231, in the consistent categories on day 2. This result indicates that not much additional learning was taking place on day 2 in the consistent categories. In line with this interpretation, in the consistent categories the average PE from the first block on day 2 was not significantly different from that at the end of day 1, *t*(24) = 1.01, *p* = .322, indicating that participants utilised the schema learned on day 1 from the beginning of day 2.

In contrast, the average PE in the first block of the inconsistent categories was significantly higher than the average PE in the last block of day 1, *t*(24) = 5.305, *p* < .001. Moreover, in the inconsistent categories there was a significant main effect of block, *F*(9, 216) = 3.520, *p* < .001, Sphericity assumed. PE decreased linearly across blocks, evident in a significant linear trend, *F*(1, 24) = 8.956, *p* = .006. Notably, however, performance in the inconsistent condition on day 2 never reached the same level as performance in the last block on day 1: A comparison of block 10 of day 1 and block 10 of day 2 in the inconsistent categories yielded a significant difference, *t*(23) = 3.15, *p* = .0045. This last finding suggests that, at the end of day 2 participants were potentially still confused by the initially learned schema in the inconsistent condition. Further evidence for this interpretation will be provided in the following sections.

#### Category consistency affects old and new schema-based responding in the Final Test on day 3

##### Modelling and Permutation Testing

I used computational modelling and permutation testing to assess the hypothesis that, in the inconsistent condition on day 3, participants would show a bias towards the location of the ‘old’ *previously* relevant schema mean, that is the mean location that a specific category occupied on day 1. Since the day 1 schema mean in the inconsistent condition was always −90 degrees from the ‘new’ day 2 schema mean, such a bias would be evident in an increased number of responses 90 degrees counter-clockwise to the current new schema mean in the inconsistent condition compared to the consistent condition. In contrast, in the consistent condition, since the schema never changed, participants had no reason to be biased towards responses 90 degrees counter-clockwise to the current schema.

To test whether there was a bias in responses on day 3, I applied a mixture model to the data collected in the final test on day 3. The model comprised (i) a von Mises distribution centred around the feedback locations, (ii) a uniform distribution component, and (iii) an additional von Mises distribution centred around −90 degree from the currently relevant ‘new’ schema mean. The von Mises distribution centred on the feedback locations captured memory for the studied location. The uniform distribution captured random guesses (∼forgotten trials). The second von Mises distribution was included to capture responses that were consistent with the ‘old’ schema mean in the inconsistent categories. This approach is similar to that of modelling ‘non-target’ responses in previous work (Bays, Catalao, & Husain, 2009). The model was run on data pooled across participants to enhance stability of the model, and modelling was done separately for consistent and inconsistent categories, as done in previous research (cf. Richter et al., 2019). Only trials originally learned on day 2 were used for this analysis, as it focuses on the effect that the change in a schema on day 2 has on subsequent memory on day 3 (cf. Richter et al., 2019).

As outlined above, I tested whether participants would show a tendency to bias their responses towards the ‘old/day 1 schema mean’, in the inconsistent categories. For the inconsistent condition, this day 1 schema mean was always presented −90 degrees from the current/new schema mean. To test my hypotheses I used an approach which is similar to the one use in a previous paper (Richter et al., 2019), and which is described further in Schneegans and Bays (2016). For each trial, I calculated the *posterior probability* that the response a participant gave stemmed from each of the three mixture components mentioned above. To allow for a valid comparison with the inconsistent categories, in the consistent categories, where the ‘old/day 1’ schema mean is identical to the current/new schema mean, I also modelled the probability of responding −90 degrees from the current schema mean. That is, −90 degree responses were modelled for both the consistent and inconsistent conditions, but in the inconsistent conditions these responses corresponded to the location of the old/day 1 schema mean.

To assess the hypothesis that there would be fewer trials assigned to the −90 degree model component in the consistent categories than the inconsistent categories, I used permutation testing. The results of this analysis indicated more responses around −90 degrees, that is in the ‘old schema mean’ location, for the inconsistent than the consistent categories, with *p* < .001 and 1000 iterations, consistent with participants still being biased by the day 1 schema in the inconsistent condition.

#### Prediction error at day 2 predicts schema updating on day 3 more strongly in the inconsistent conditions

The modelling analysis suggests that, on day 3, participants showed a stronger tendency to recreate locations −90 degrees from the current/new schema in the inconsistent vs. consistent conditions. In other words, the preceding analysis showed that, on day 3, participants are biased towards the ‘old’ schema mean in the inconsistent conditions. In a next step I asked whether this day-3 bias is *diminished* for trials in which larger PEs occurred on day 2. Such a finding would provide support for the central hypothesis that PEs might *trigger* schema updating. In other words, if PEs are relatively large on day 2, does this induce more learning/updating of the schema, evident in a smaller bias on day 3?

##### Schema-updating score

To measure “schema updating” I calculated the degree to which participants responded consistent with the ‘new’ schema on day 2 and day 3. To assess schema-based responding, I calculated the absolute difference between the responses given by the participants and the new *schema mean* (i.e., an error measure between response and schema mean, rather than the traditional error measure between response and the location learned in the feedback phase) on day 2 and day 3 respectively. Subsequently, I calculated a *schema updating score* for each trial by subtracting these day 3 final test error values from the corresponding values during new learning on day 2. To make error values comparable between both sessions, errors were *z*-scored first. As day 3 errors were subtracted from day 2 errors, *positive schema-updating scores* indicate that responses are closer to the current/new schema mean in the final test (day 3) than in the new learning phase (day 2), indicating a stronger orientation towards the current/new schema, and hence *more updating*.

To assess whether PEs predict the degree to which updating towards the current/new schema occurs, I correlated the PEs at day 2 with the calculated schema updating scores. I then compared the resulting mean correlations between the consistent and inconsistent categories across participants. I found, as expected, that there was a positive mean correlation between the updating measure and the absolute PE. This effect was found for both consistent, *r*_*consistent*_ *=* .27, *t*(1,24) = 6.445, *p* < .001, and inconsistent categories, *r*_*inconsistent*_ *=* .39, *t*(1,24) = 24.740, *p* < .001. This outcome indicates that in either condition larger PEs during NL resulted in learning. Supporting my hypothesis, this correlation was stronger in the inconsistent than in the consistent categories, where more learning was expected to take place, due to the change in the schema mean, *t*(1,24) = 3.260, *p =* .0033, Figure 4A. It follows that larger PEs induced learning in general, but this effect was most pronounced when a schema needed to be updated.

##### Modelling

A second way to assess the relationship between PEs on day 2 and memory on day 3 is to examine the relationship between PE and the model derived-probability of responding at −90 degrees. To recap, these −90 degree responses capture responding with the ‘old/day 1’ schema mean in the inconsistent categories. For the consistent categories, −90 degree responses provide a baseline of the probability of responding at this location that would be expected if the schema is never updated.

Turning to the consistent categories first, there was a significant relationship between the size of PEs and the probability of errors around −90 degrees, *r*_*consistent*_ *=* .145, *t*(1,24) = 3.58, *p* < .002. The result of this analysis is presented in Figure 4B. As can be seen from the figure, large PEs in the consistent categories were associated with an *increased* probability to respond around −90 degrees. This result may seem surprising at first, as it indicates that large PEs on day 2 lead to relatively large errors during responding at the final test. However, as participants generally committed low errors in the consistent categories on day 2, large PEs on day 2 were observed primarily for trials further away from the schema mean. These trials that occurred on the ‘outer boundaries’ of the category’s 90-degree circle segment naturally led to larger errors (see Richter et al., 2019 for similar results, evidencing more schematic responding in the consistent category). At the same time, these trials would not induce ‘updating’ of the schema, as they would be perceived as expected noise, based on the same noise level experienced on day 1, thus explaining the observed relationship.

The data from the consistent categories therefore provides an important baseline for the relationship between PEs and number of −90 degree responses expected if the schema never changes. Consequently, the next goal is to determine whether larger PEs in the inconsistent categories, resulted in a *relative decrease* in the probability to respond with the old schema mean compared to the consistent categories, or in other words, if the observed relationship between PEs and −90 degree responses is *weakened* for the inconsistent categories.

Consistent with this prediction, larger PEs on day 2 were less strongly associated with −90 degree responses on day 3 in the inconsistent than the consistent categories, *t*(1,24) = −2.802, *p =* .010. Specifically, in the inconsistent categories, larger PEs predicted −90 degree (or ‘old’ schema mean) responses only numerically, *r*_*inconsistent*_ *=* .05, *t*(1,24) = 1.684, *p* = .110, suggesting that larger PEs may have led to an updating of the schema, at least in some trials. This finding is particularly noteworthy since, in the inconsistent but not the consistent categories, participants had been presented with items at the −90 degree position, that is the ‘old’ schema mean, on day 1. Still, large PEs on day 2 were not predictive of responses guided by the old schema in the inconsistent compared to the consistent condition. This finding is therefore again consistent with the aforementioned hypothesis that following larger PEs more updating took place in the inconsistent condition. In sum, both the schema-updating analysis and the modelling results suggests that larger PEs on day 2 result in more updating of schema inconsistent information on day 3.

#### Confidence is negatively related to PE

Participants’ confidence ratings were also collected in order to investigate how confident participants were in their location responses on day 2. A correlation was calculated between the confidence rating a participant gave on each trial and the subsequently observed absolute PE. A tendency for a negative relationship (lower PE for high confidence responses) indicating that participants had the metacognitive ability assess their performance was observed in both the consistent and inconsistent condition. However, this effect was only significant for the consistent categories, mean *r*_*consistent*_ = -.0619, *t*(24) = −2.5564, *p* = .0173, but not the inconsistent categories, mean *r*_*inconsistent*_ = -.0245, *t*(24) = −0.9998, *p* = .3274. However, as the difference of correlations between the consistent and inconsistent categories was not significant, *t*(24) = 1.1087, *p* = .2785, these results will require a cautious interpretation.

#### Interim summary of behavioural results

To summarize the behavioural results before considering the EEG analysis, the data indicates that participants successfully learned the schema on day 1. For the inconsistent categories they updated their schemas on day 2, but still remained below day 1 performance (Figure 4A). On day 3 participants showed a bias towards responses corresponding to the old schema mean in the inconsistent categories. However, the size of the PE on day 2 predicted a reduction in this bias, evident in schema updating on day 3. This relationship between PE and updating was stronger in the inconsistent condition (Figure 4B). Across trials high confidence was generally associated with lower PEs, but this effect was reliable only in the consistent condition (Figure 4B).

### EEG Results

Data was analysed both using grand-averages as well as single-trial data. For grand-average analyses data was not filtered. To assess the effect of condition (consistent versus inconsistent categories) on P3 amplitude a 100 ms time window 350-450 ms after the feedback stimulus was used. For analyses focussing on single-trial data, data was first filtered using a 6Hz filter. To reduce the effect of outliers for the single trial analyses, the 2.5% highest and lowest amplitudes were excluded from the analyses retaining only the central 95% of trials. This procedure was done separately for the consistent and inconsistent conditions, to ensure that equal trial numbers were excluded in both conditions. For the trial-wise analysis the mean amplitude in the same P3 window as stated above was used. For any across-subject analyses, single-trial P3 amplitudes and mean absolute PEs were furthermore z-scored, to account for between subject differences. As noted above, for schema-updating scores z-scoring was already implemented during calculation of these values and schema-updating scores were accordingly not z-scored a second time.

#### Larger P3s for the inconsistent categories

The EEG analysis focused on two clusters of interest: fronto-central and centro-parietal electrodes. These locations have shown to display belief-updating effects (Bennett et al., 2015; Jepma et al., 2018, 2016; Kolossa et al., 2015) in previous studies and have displayed involvement in subsequent memory effects in studies of long-term memory (e.g., Richter & Yeung, 2016). I first tested the prediction that the P3 should be more pronounced in the inconsistent condition rather than in the consistent condition during day 2 of learning. Supporting this prediction, grand-average results showed that in the frontal electrode cluster trials of the inconsistent categories evoked a larger P3 response than trials of the consistent categories, *t*(24) = −2.553, *p* = .018. A similar, marginally significant effect was observed in the parietal sites, *t*(24) = −2.063, *p* = .050. The difference in the size of effects on frontal and parietal sites was not significant, *t*(24) = 1.197, *p* = .243.

**Figure 5.**
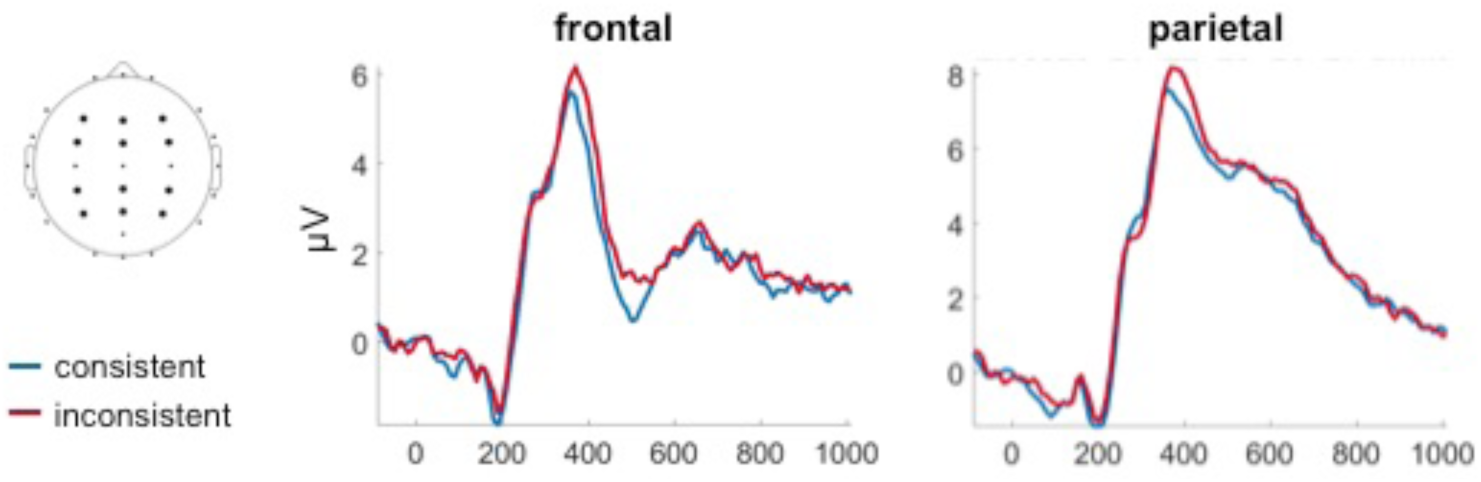
Grand-average feedback-evoked P3 at frontal and parietal sites. The average signal was calculated across clusters of 6 electrodes displayed as black circles in the topography on the left. EEG amplitude is plotted separately for the consistent (blue) and inconsistent (red) categories.

#### PE predicts trial-by-trial P3 amplitude

I examined whether P3 amplitude increased reliably with increasing PE at frontal and posterior sites. At the frontal cluster increasing PE size was associated with increasing P3 amplitude (Figure 6A). This effect was significant when computed across all subjects and trials, *t* = 2.85, *p* = .0044, and also when computed within subjects, *t*(24) = 2.68, *p* = .013, in the inconsistent categories. A corresponding effect was not observed in the consistent categories, both when computed across all subjects and all trials, *t* = 0.279, *p* = .783, as well as when assessed within subjects, *t*(24) = 0.187, *p* = .852. The difference in the within-subject P3-PE correlation between consistent and inconsistent categories was marginally significant, *t*(24) = 1.85, *p* = .075.

**Figure 6.**
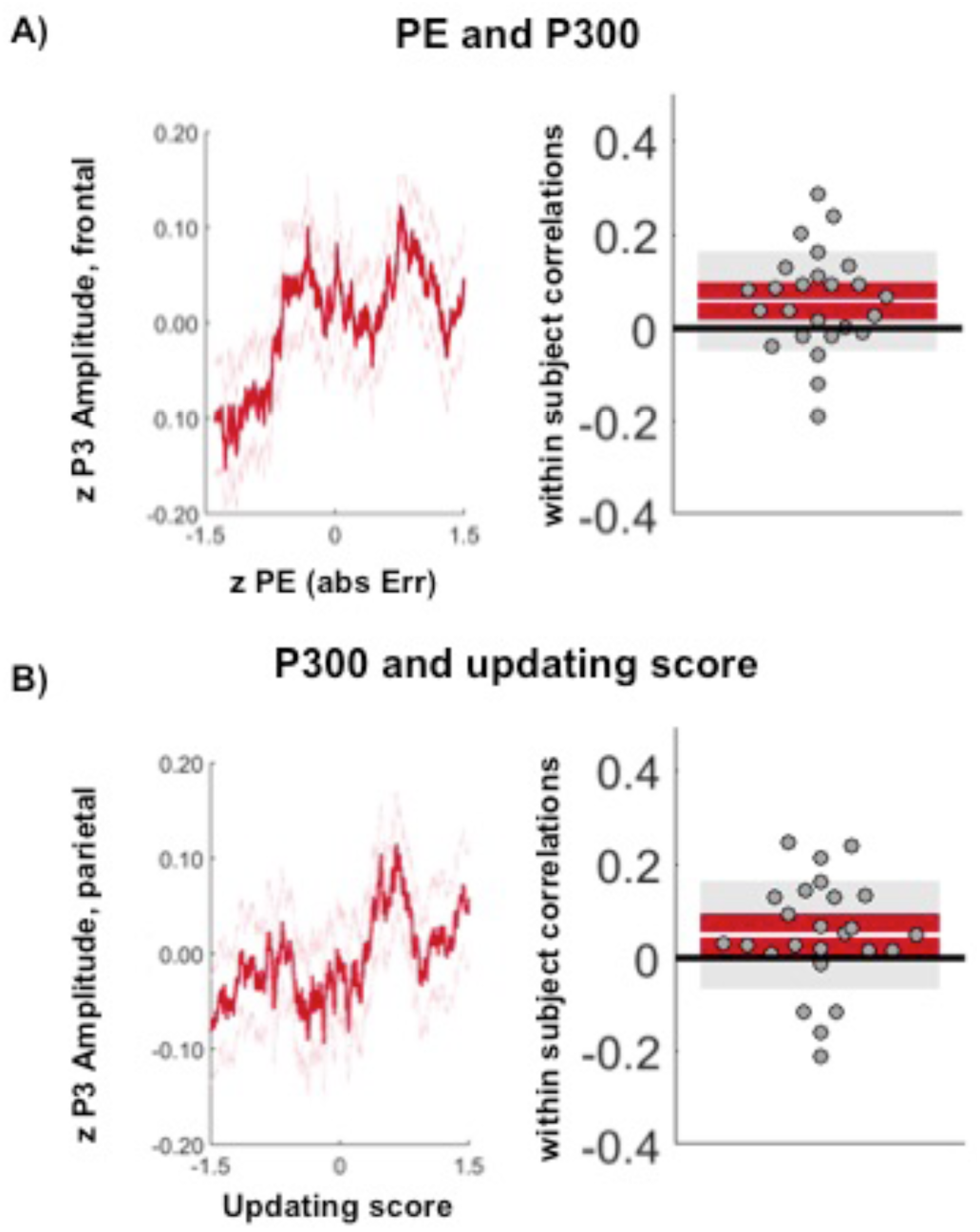
Relationship between PE and P3 amplitude as well as between P3 amplitude and updating scores in the inconsistent condition. No significant effects were found for the consistent condition. In the left panels data is pooled across participants. PE and updating score are averaged across running bins of 300 trials for display purposes, but statistics reported in the text are conducted on trial-wise data. In the right panels each circle represents one subject. The white line represents the mean, and the red area the 95% confidence interval. The grey areas indicate *mean* ± 1 *SD*. Illustration of the within-subject correlation was done via the notBoxPlot function in Matlab https://nl.mathworks.com/matlabcentral/fileexchange/26508-notboxplot. (**A**) Relationship between PE and P3 amplitude at frontal electrodes. (**B**) Relationship between P3 amplitude and updating of the schemas in the final test on day 3 at parietal electrodes. Schema updating was calculated as the difference in absolute error from the schema mean between new learning on day 2 and final test on day 3 (day 2 - day 3; accordingly positive values mean updating has taken place, see details in text).

For the posterior cluster, numerical effects in a similar direction were observed. However, they were not statistically reliable (inconsistent condition: across all subjects and trials, *t* = 0.707, *p* = .486; within subjects, *t*(24) = 0.495, *p* = .620; consistent condition: across all subjects and trials, *t* = 0.108, *p* = .915; within subjects, *t*(24) = 0.039, *p* = .969), and there was no significant difference between the inconsistent and consistent categories in the within-subject effects, *t*(24) = 0.346, *p* = .733.

To assess location differences, I compared whether the mean within-subject P3-PE correlation was stronger at frontal than posterior electrodes. The average within-subject correlation was marginally stronger at frontal than posterior electrodes in the inconsistent condition, *t*(24) = 1.790, *p* = .086. Consistent with an overall lack of an effect of PE on P3 amplitude in the consistent condition, there was no difference to be observed between frontal and posterior electrode sites, *t*(24) = 0.1383, *p* = .8912.

Thus, PE predicted P3 size and this effect was marginally stronger in the inconsistent than the consistent categories and at frontal rather than posterior electrode sites.

#### Trial-by trial P3 amplitude predicts schema updating

If schema updating is initiated by the P3, P3 should not only correlate with the PE, but also with the degree to which participants updated their responses on day 3. To test this prediction, I correlated P3 amplitude in the consistent and inconsistent condition with the schema-updating score (see calculation above). Analyses revealed that in the inconsistent condition there was a positive correlation between P3 amplitude and later schema updating. This relationship was marginal at frontal electrodes when investigated across all subjects and trials, *t* = 1.90, *p* = .057, as well as within subjects, *t*(24) = 1.88, *p* = .071. For the consistent categories, the relationship was insignificant both across all subjects and trials, *t* = 1.135, *p* = .256, and within subjects, *t*(24) = 0.982, *p* = .336. However, there was no significant difference between the within-subject effects of the consistent and inconsistent categories, *t*(24) = 0.362, *p* = .721.

At posterior electrodes (see Figure 6B), a significant positive relationship between P3 amplitude and updating was found for the inconsistent categories, again across all trials, *t* = 2.41, *p* = .016, as well as within subjects, *t*(24) = 2.13, *p* = .044. The same analysis for the consistent categories revealed an insignificant effect across all trials and subjects, *t* = 1.44, *p* = .151, and a marginal effect within subjects, *t*(24) = 1.94, *p* = .065. Again, there was no significant difference between the within-subject effects of the consistent and inconsistent categories, *t*(24) = 0.651, *p* = .522.

To assess location differences, I compared the mean within-subject correlation for each location by category consistency combination for the relationship between P3 and schema-updating score. Again, no difference between frontal and posterior sites in the average within-subject correlation was observed neither for the inconsistent, *t*(24) = −0.4943, *p* = .626, nor the consistent condition, *t*(24) = −0.316, *p* = .755.

The data therefore revealed that P3 amplitude predicted schema-updating and this effect was significant in the inconsistent condition at posterior electrode sites. Effects were observed in a similar direction at frontal electrode sites and for the consistent condition, but they did not reach significance.

I also investigated whether larger P3s were associated with a decreased probability of responding with the ‘old’ schema mean (derived from the model, see calculation above). While the effect was numerically in the predicted direction – with a negative correlation indicating fewer ‘old’ schema mean responses for larger P3s – it did not reach significance for either the consistent or inconsistent categories both a frontal or at posterior electrodes in both the within- and across-subject analyses (all *ps >* .4352).

To summarize the results of the EEG data, the inconsistent categories resulted in larger P3s, and P3 amplitude correlated with behavioral PEs, especially at frontal electrodes. P3 amplitude also correlated with schema-updating scores, but this effect seemed to be reliable only at posterior electrode sites. Overall, it follows that P3 amplitude tracked both, PEs and schema-updating.

## Discussion

The current experiment tested the hypothesis that similar neural mechanisms underlie the updating of long-term memory schemas as do the updating of short-lived beliefs in decision-making contexts. For this purpose, I used a continuous report memory task, in which participants predicted the location of stimuli belonging to different categories in a three-day study. By manipulating the consistency of the categories on day 2, after a 24-hour consolidation phase, I induced PEs in participants. PEs were predictive of behavioural updating performance as well as P3a size, while P3b amplitude additionally correlated with schema-updating, revealing that similar neural mechanisms may underlie the flexible modification of complex knowledge structures and short-lived beliefs.

### Relationship to neural processes underlying belief updating

The neural mechanisms of schema updating have so far remained elusive, and to date have only been speculated upon (e.g., Richter et al., 2019; van Kesteren et al., 2012). Previous research that has studied the link between trial-by-trial variations of prediction violations and updating performance has largely focused on much shorter time scales and assessed trial-by-trial predictions typically within only one testing session (e.g., Bennett et al., 2015; Jepma et al., 2018, 2016; Kolossa et al., 2015). In contrast to these previous studies, the current experiment does not focus on neural correlates of the updating of *immediate* predictions in the task, but rather updating effects evident in the participants’ performance *one day later*, in a final memory test. My data strongly suggests that similar neural mechanisms underlie PE-based updating of long-term memory schemas, as they do short-term beliefs (Bennett et al., 2015; Jepma et al., 2018, 2016; Kolossa et al., 2015). Specifically, the current study tracked changes to these complex mnemonic structures across days and linked these changes to the P3 signal.

Consistent with a role of the P3a in mismatch detection and novelty processing, I found that PE correlated with P3a size. As mentioned above, P3a has been linked to hippocampal processing. Recent findings suggest that interactions between hippocampus and the dopaminergic ventral tegmental area play a crucial role in learning based on PE in declarative memory (Calderon et al., 2020). The correlation between PE and P3a size in the current data might reflect these processes and consequently PE-induced dopaminergic learning mechanisms.

When investigating the relationship between P3 amplitude and the degree to which participants updated their responses in the final test on day 2, I found that P3b amplitude significantly predicted updating in the inconsistent categories. The P3b signal has been associated with contextual updating or more broadly with the maintenance of relevant representational context (Polich, 2007, 2012). Similarly, in the current study, the P3b could index an update to the current context: the new, shifted schema. In other words, the correlation between updating and the P3b signal could reflect neural processes that allow modification of a now out-dated model of the world.

Looking at the relationship between both components, there were no significant differences between P3a and P3b effects. For this reason, the exact involvement of P3a and P3b in prediction violations and in the subsequent updating of schemas needs to be further investigated. It seems likely that both effects operate in conjunction to update memory schemas based on prediction violations, possibly reflecting the involvement of distributed brain networks (see below).

### Updating of complex knowledge structures

The current study perhaps differs most notably from previous work with regards to the kind of information that needs to be updated. Above, a distinction to belief-updating tasks has already been drawn. While PEs are traditionally studied in the context of implicit learning, PEs have recently also been related to declarative learning (see Ergo, De Loof, & Verguts, 2020, for a review). Previous studies on the link of declarative learning and PEs have focused on single associations, episodic memories, or semantic facts (De Loof et al., 2018; Pine et al., 2018; Rouhani, Norman, & Niv, 2018). The current work brings a qualitatively different kind of declarative memory into the focus by studying more complex knowledge structures, rather than individual mnemonic entities. Moreover, it does not investigate mnemonic performance per se (i.e., general encoding success), but instead differences in the *updating or modification* of higher-level memory constructs.

Previous research has only speculated that such updating of knowledge structures such as schemas might rely on a mismatch between a schema-based prediction and new encountered information (Richter et al., 2019; van Kesteren et al., 2012), since a correlation between PE and learning has been observed for other types of memory (Greve et al., 2019; Henson & Gagnepain, 2010; Krawczyk, Fernández, Pedreira, & Boccia, 2017; Pine et al., 2018). However, due to the absence of sufficiently sensitive methods to measure trial-wise differences in schema updating in the laboratory, it has been difficult to identify the source of gradual changes to complex memory structures such as schemas.

The continuous report task used in the current paradigm delivered the essential tool to induce trial-by-trial differences in PE size, providing me with a powerful method to directly relate trial-wise differences in PE and P3 size to schema updating. The observed relationship between PEs and schema-updating extends prior findings of a link between prediction violations and learning (Greve et al., 2017, 2019; Pine et al., 2018; Sinclair & Barense, 2018, 2019) to the flexible updating of complex knowledge constructs. This finding therefore provides unique insights into how integrated and abstracted dynamic knowledge constructs are maintaining their flexibility.

### Non-reward based prediction errors

The vast majority of studies on learning based on prediction violations has used reward-prediction paradigms in which participants encode information while varying levels of reward are given. In these studies the PE is therefore based on the size and expectancy of externally provided reward rather than the stimuli themselves. The current study differs from this previous work as no direct reward is given.

A small number of recent studies have also investigated non-reward based learning. For example, studies have based PEs on participants’ subjective confidence ratings (e.g., Butterfield & Metcalfe, 2001). These studies have found that high confidence errors lead to more learning than low confidence errors (Metcalfe, 2017). In contrast to this previous work, in the current study the link between confidence and PE was only reliable in the *consistent* condition, even though a stronger link between PEs and updating was observed in the *inconsistent* condition. This finding that the correlation between confidence and PE was only significant for the consistent categories suggests that this relationship is evident in situations in which the participants can be relatively certain about their answers. In the current study participants may have experienced overall reduced certainty in their responses in the inconsistent condition. Confidence judgments might not be an ideal measure of prediction violations in changing or highly uncertain environments like this, because a participant’s model of the world is in the process of changing. In such a situation confidence judgements may not adequately track trial-by-trial performance. The role of uncertainty and confidence in schema-updating will therefore have to be further assessed in future work. Confidence may be most useful if identical information is probed repeatedly, because it closely tracks objective performance in this case. It may be less suitable if new predictions are made based on the gradual abstraction of information in a noisy environment. In this context the employed memory precision measures are of advantage, because they provide us with an objective and continous (rather than categorical) measure of performance that can be related to neural effects as well as later updating, and that seems to be potentially less affected by uncertainty.

### Network mechanisms of schema updating

Regarding the neural mechanisms underlying PE-based updating of memory schemas, likely a large network of brain areas and neurotransmitter systems is involved. The P3 is a signal that has been closely linked to catecholaminergic mechanisms in the brain (Polich, 2012). A distinction has been made between the neurotransmitter systems involved in the P3a and P3b. The more frontal P3a has been linked to dopaminergic activity (Polich, 2007, 2012; Polich & Criado, 2006) and recent research has demonstrated that reward-related PEs, previously strongly linked to the dopamine system (Schultz, Dayan, & Montague, 1997), are positively correlated with successful memory encoding (Greve et al., 2017; Jang, Nassar, Dillon, & Frank, 2019; Rouhani et al., 2018). It follows that learning based on prediction violations in schema memory might share neural mechanisms with that of reward-mediated episodic memory. Consequently, external reward may not be necessary in updating long-term beliefs. In fact, it has been argued that good performance is in itself rewarding (Satterthwaite et al., 2012). The rewarding effect of performing well might therefore be a significant motivator in the current study, and could explain the involvement of similar neural mechanisms in the current task and in paradigms studying reward-based learning.

The neural processes underlying the P3b may rely on somewhat distinct neurotransmitter systems than those of the P3a: In addition to being affected by dopamine (Polich, 2007; Polich & Criado, 2006), the P3b potential has also been linked to the locus coeruleus norepinephrine system (De Taeye et al., 2014; Jepma et al., 2018, 2016) and has been suggested to index target-detection, response selection, or rate of learning in the context of decision-making, especially when the environment becomes uncertain unexpectedly (Yu & Dayan, 2005). The relationship between P3b and updating in the current study could therefore index increased learning due to unexpected uncertainty in the inconsistent condition. After participants discovered regularities amongst the noise in the schemas on day 1, a subset of the schemas changed without an obvious reason on day 2, and the resulting uncertainty induced learning, or in other words *schema updating*.

How do we combine the findings on P3a and P3b? While the P3a is believed to be generated by frontal areas (Ebmeier et al., 1995; Kirino, Belger, Goldman-Rakic, & McCarthy, 2000), the P3b has been suggested to be generated by temporo-parietal regions (Polich, 2012). The involvement of both frontal and posterior ERP components in the current study is therefore consistent with the idea proposed in the literature that schema updating might be dependent upon two different different networks. The initial identification of schema incongruent information may be subserved by fronto-hippocampal interactions, most likely involving (ventro-) medial prefrontal cortex (van Kesteren et al., 2012). Subsequent updating may recruit a more posterior hippocampal temporo-parietal network (cf. van Kesteren et al., 2012). Future research that can more accurately track the localisation of these effects is needed to investigate the regionally specific brain mechanisms underlying schema updating. It will be of particular interest to understand how independent the two proposed mechanisms – the top-down attention capture by a stimulus in the inconsistent condition, and the subsequent updating of the newly encoded information into the pre-existing memory schema – operate.

## Conclusion

Together, the current data provide novel insights into how complex memory schemas remain flexible in changing environmental conditions. With regards to the relevance of such knowledge structures in many areas of cognition such as education (Bransford, Brown, & Cocking, 2000; Ruiter, van Kesteren, & Fernandez, 2012) or decision making (Gilboa & Marlatte, 2017; Hebscher & Gilboa, 2016), understanding the mechanisms that govern this kind of learning is of critical importance. The current paper provides evidence that memory schemas, similar to more short-lived belief structures, are updated via prediction-error based learning, and that similar neural mechanisms, indexed by the P3, underlie updating in both cases. Moreover, this updating can occur over larger time scales than what has been tested before - at least 24 hours - and is not limited to the updating of immediate behaviour.

## Acknowledgements

I would like to thank Angélica van der Ploeg and Alexandra Felea for help with data collection.

